# Intra-annual temperature variability and nutrient enrichment drive seasonal β-diversity in global grasslands

**DOI:** 10.1101/2022.10.24.513509

**Authors:** Magda Garbowski, Elizabeth Boughton, Anne Ebeling, Philip Fay, Yann Hautier, Hanna Holz, Anke Jentsch, Stephanie Jurburg, Emma Ladouceur, Jason Martina, Timothy Ohlert, Xavier Raynaud, Christiane Roscher, Grégory Sonnier, Pedro Maximiliano Tognetti, Laura Yahdjian, Peter Wilfahrt, Stan Harpole

**Author notes:** **Corresponding author email:**, Corresponding author’s present address: Botany Department, 1000 E. University Ave. University of Wyoming, Laramie, WY 82071. **Open Research Statement:** Estimates of observed, expected, and null seasonal temporal diversity used in these analyses as well as the novel code used to generate these estimates can be found at: https://github.com/MagdaGarbowski/Nutrient_Network_Seasonality. Upon acceptance, these data and finalized code will be publicly available and archived on figshare.

## Abstract

In many grasslands, species with specific traits occupy unique temporal positions within communities. Such intra-annual segregation is predicted to be greatest in systems with high intra-annual climate variability because fluctuating environmental conditions provide opportunities for temporal niche partitioning among species. However, because most studies on intra-annual community dynamics have been conducted at individual sites, relationships between intra-annual climate variability and seasonal community dynamics at global scales have not yet been identified. Furthermore, the same characteristics that promote species-specific responses to fluctuations in environmental conditions may also drive species-specific responses to global change drivers such as eutrophication. Research provides evidence that eutrophication alters inter-annual plant community dynamics yet understanding of how it alters intra-annual dynamics remains limited.

We used early-season and late-season compositional data collected from 10 grassland sites around the world to ask how intra-annual variability in precipitation and temperature as well as nutrient enrichment shape intra-annual species segregation, or seasonal β-diversity, in plant communities. We also assessed whether changes in the abundances of specific functional groups including annual forbs, perennial forbs, C3 and C4 graminoids, and legumes underpin compositional differences between early- and late-season communities and treatments. We found that intra-annual temperature variability and seasonal β-diversity were positively related but observed no relationship between intra-annual precipitation variability and seasonal β-diversity. This suggests that positive relationships between α-diversity and intra-annual temperature variability identified in earlier studies may be underpinned by the positive influence of intra-annual temperature variability on temporal segregation of species within growing seasons. We found that nutrient enrichment increased seasonal β-diversity via increased turnover of species between early- and late-season communities. This finding mirrors patterns observed at inter-annual scales and suggests fertilization can alter compositional dynamics via similar mechanisms at varied temporal scales. Finally, fertilization reduced the abundance of C4 graminoids and legumes and eliminated intra-annual differences in these groups. In contrast, fertilization resulted in intra-annual differences in C3 graminoids which were not observed in control conditions, and increased abundance of C3 graminoids and annual forbs overall. Our study provides new insight into how intra-annual climate variability and nutrient enrichment influence biodiversity and seasonal dynamics in global grasslands.

## Introduction

Ecologists have investigated the mechanisms by which climate shapes plant diversity for decades (e.g., Rosenzweig 1995, Hawkins et al. 2003, Harrison et al. 2020). While most research examines associations with mean climate conditions, numerous theoretical (e.g., Lewontin & Cohen 1969, Chesson 2000, Adler & Drake 2008) and empirical (e.g., Levine & Reese 2004, Letten et al. 2013) studies have highlighted the importance of temporal climate variability in structuring populations and communities. Central to these studies is the notion that environmental fluctuations provide opportunities for temporal niche partitioning among species (e.g., Chesson & Huntly 1997, Chesson 2000). Intra-annual fluctuations in temperature and precipitation may be particularly important drivers of intra-annual temporal niche partitioning in plant communities as they influence plant physiology and phenology and modulate ecosystem processes related to water availability and nutrient cycling (Luo et al. 2020). Modeling studies have demonstrated that intra-annual variation in precipitation supports coexistence among plant species with slight ecological differences in germination requirements, growing season length, or seasonal growth activity (Mathias and Chesson 2013). Empirical evidence documenting the influence of intra-annual climate variability on temporal niche segregation mainly comes from studies of desert systems in which winter and summer annuals segregate temporally in response to seasonal fluctuations in precipitation (e.g., Mulroy & Rundel 1977, Guo & Brown 1997). But temporal segregation of coexisting species is evident in diverse ecosystems including tropical forests (Sapijanskas et al. 2014) and grasslands (Fargione & Tilman 2005).

In many grasslands, species with specific strategies optimally utilize resources under different environmental conditions and thus coexist by occupying unique temporal positions within communities. For example, in grasslands co-dominated by C3 and C4 grasses, C3 grasses grow and set seed primarily in early to mid-season and C4 grasses grow and set seed primarily in late season. Similar dynamics occur in communities without C3 and C4 grasses present. For instance, in temperate European grasslands early-season grasses and forbs contribute most to productivity at the beginning of the season, whereas late-season grasses contribute most to productivity late in the growing season (e.g., Guimarães-Steinicke et al. 2019, Doležal et al. 2017). However, the same characteristics that promote species-specific responses to fluctuations in environmental conditions often drive species-specific responses to global change drivers such as eutrophication. For example, subordinate forbs and legumes that thrive at different time periods from dominant grasses are often reduced in fertilized conditions in European meadows (e.g., Doležal et al. 2017). Research from the last several years has elucidated how eutrophication alters inter-annual plant community dynamics (e.g., Hautier et al. 2014, Chen et al. 2021). Yet, interactions between global change drivers and intra-annual dynamics of plant communities remain largely unexplored (White & Hastings 2020).

Eutrophication in grasslands leads to reduced plant species richness and diversity (e.g., Hautier et al. 2009, Harpole et al. 2016). Evidence for these patterns has primarily been gathered from studies that measure plant community composition once a year, usually at the peak of biomass production (e.g., Borer et al. 2014). However, research has demonstrated that diverse resource-use strategies allow distinct species assemblages to dominate in different points within a growing season (Doležal et al. 2017, Guimarães-Steinicke et al. 2019, Huang et al. 2019). In grassland systems, nutrient enrichment may promote the dominance of species with specific traits for longer periods of time and reduce overall dissimilarity among early- and late-season communities (Doležal et al. 2017). Further, the addition of multiple limiting nutrients may reduce the importance of species-specific trade-offs associated with competition for particular nutrients (Harpole et al. 2016) that are most limiting at different points throughout the growing season (Klaus et al. 2016). Assessing how nutrient enrichment affects compositional dissimilarity between early and late-season assemblages would clarify whether patterns of reduced diversity in response to fertilization at inter-annual scales are underpinned by reduced dissimilarity at intra-annual scales.

Changes in overall diversity in responses to nutrient enrichment often result from changes in the abundance of species that are affected by nutrient enrichment in different ways. Nitrogen enrichment reduces the abundance of legumes and nutrient conservative C4 grasses but increases the abundance of nutrient acquisitive C3 grasses (e.g., Suding et al. 2005, Tognetti et al. 2021). Fertilization with multiple nutrients can further alter community composition by causing shifts in the relative abundance of species from specific functional groups (Wilcots et al. 2021). When nutrient enriched communities become dominated by specific species or functional groups, ecosystem stability may be reduced via a loss of asynchronous species responses to environmental fluctuations (Hector et al. 2010, Hautier et al. 2014). In addition, nutrient additions that result in the loss of forb and legume species can have cascading effects on organisms at other trophic levels, such as pollinators, that often depend on these groups for floral resources at specific times of the year (Burkle & Irwin et al. 2010, Dyer et al. 2021). Understanding how nutrient enrichment affects the abundances of specific functional groups at specific timepoints in the growing season (i.e., early vs late) and between timepoints is essential to identifying when compositional changes take place and which species drive them.

In this study, we used above-ground species composition data collected early and late in the growing season from 10 grassland sites around the world to assess how intra-annual variability in temperature and precipitation as well as eutrophication influence early vs. late season dissimilarly (seasonal β-diversity hereafter) of plant assemblages. The 10 sites are part of the Nutrient Network – a globally replicated experiment in which herbaceous plant communities are supplemented with fertilizer. Data from each site spanned between 4 and 11 years. We used data from untreated plots to assess relationships between intra-annual precipitation and temperature variability (estimates obtained from WorldClim; Fick & Hijmans 2017) and seasonal β-diversity across our study sites. We then examined the effects of fertilization on seasonal β-diversity and its components of nestedness and replacement. Finally, to determine which species underpin changes in seasonal β-diversity in response to fertilization, we assessed overall changes in abundance of different functional groups (i.e., C3 graminoids, C4 graminoids, annual forbs, perennial forbs, legumes) between early and late timepoints and fertilization treatments. Our hypotheses were that:

H1. Sites with high intra-annual variability in temperature and precipitation would have high seasonal β-diversity.
H2. Fertilization would decrease seasonal β-diversity because a subset of species would dominate across early and late sampling timepoints (i.e., fertilized plots would have higher nestedness).
H3. Differences in early-season vs. late-season community composition in response to fertilization would be underpinned by increased abundances of resource acquisitive species (e.g., C3 graminoids, annual forbs) and decreased abundances of species with more conservative strategies (e.g., C4 graminoids, perennial forbs, legumes) within and between specific timepoints (i.e., early vs. late)

## Methods

### Study design and site locations

Data for this study were collected from 10 grassland sites from around the world that are part of the Nutrient Network – a globally distributed experiment in which plant communities are supplemented with factorial combinations of nitrogen (N), phosphorous (P), potassium (K) and micronutrients (μ) (Borer et al. 2014). While most Nutrient Network sites collect compositional data once a year, data from this subset of sites is collected twice each growing season (“early” and “late” hereafter). These sites are distributed across five continents, in Africa (ukul.za), Australia (burrawan.au), Europe (bayr.de, cereep.fr, frue.ch, jena.de), North America (arch.us, temple.us, sevi.us) and South America (chilcas.ar) (Table S1).

Compositional data used in this study were collected from 3 to 5 replicated control and NPKμ plots at these sites from between 4 (bayr.de) and 11 years (ukul.za) (mean length of experiment is 6.5 years, details in Table S1). Fertilized plots at all sites besides cereep.fr received 10 g m^−2^ of N, P, and K annually with a one-time addition of micronutrients; fertilized plots at the cereep.fr site received 2.5 g m^−2^ annually with a one-time addition of micronutrients. Despite this different application rate, all results were qualitatively similar whether the cereep.fr site was or was not included in analyses, so we retain it in the results presented here. For additional details about experimental design, please see Borer et al. (2014).

### Quantifying intra-annual temperature and precipitation variability

Estimates of intra-annual temperature and precipitation variability for each site were obtained from WorldClim (Fick and Hijmans 2017). Intra-annual temperature and precipitation variability for each site are based on monthly totals of precipitation and temperature where “Temperature variability” is the standard deviation of monthly temperature * 100 and “Precipitation variability” is the coefficient of variation of monthly precipitation totals for a given year. In both cases, higher values indicate higher variance or greater fluctuations in temperature and precipitation within a year. We chose to use annual variability values rather than growing season values because winter precipitation at some of our sites (e.g., sevi.us) is an important driver of plant community dynamics. Between 18 and 30 years of precipitation and temperature data were used to obtain average intra-annual variability values for each site based on data availability in WorldClim.

### Quantifying seasonal β-diversity

Dissimilarity indices are useful tools for measuring differences between communities in space or time. However, dissimilarity indices can be influenced by local community size (i.e., *α*-diversity) and overall richness of species at regional scales (i.e., *γ*-diversity) (e.g., Chase & Myers 2011). Deviations from null expectations of dissimilarity (e.g., z-scores) can help determine whether compositional differences, independent of local community size and regional species pools, drive dissimilarity patterns (e.g., Chase & Myers 2011). More specifically, larger z-scores indicate higher dissimilarity between communities, whereas lower deviations indicate lower dissimilarity.

To quantify dissimilarity between early and late communities in control and NPKμ treatments at each site we used deviations from expected values of the Bray-Curtis dissimilarity index (Figure 1). We did this by first permuting species abundances (i.e., absolute values of species cover) from permanent m^2^ survey plots 100 times while holding overall richness and total abundances of species constant within a plot at a given sampling timepoint (i.e., early or late season). We used 100 permutations for “early” communities and 100 permutations for “late” communities to obtain mean and standard deviation values for expected (i.e., null) dissimilarity in each plot. We then calculated z-scores for each early-to-late comparison for each plot, year, and treatment combination using the following formula:

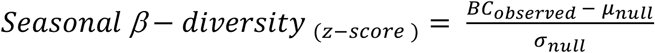

where *BC_observed_* is the observed Bray-Curtis value from each plot in each year, *μ_null_* is the expected mean for each plot in each year obtained from the distribution of permutation dissimilarly estimates, and *σ_null_* is the variance for each plot in each year obtained from the distribution of permutation dissimilarly estimates (Figure 1b). We used these z-scores as our estimates of seasonal β-diversity from each plot in each year. Comparisons from a given plot in a given year with less than four species at either the early or late sampling time point were removed prior to all analyses because communities with less than four species could not produce sufficient permutations to calculate reliable estimates of seasonal β-diversity. This resulted in the exclusion of 53 early-to-late comparisons from 888 comparisons total (i.e., 835 comparisons were included in analyses).

**Figure 1:**
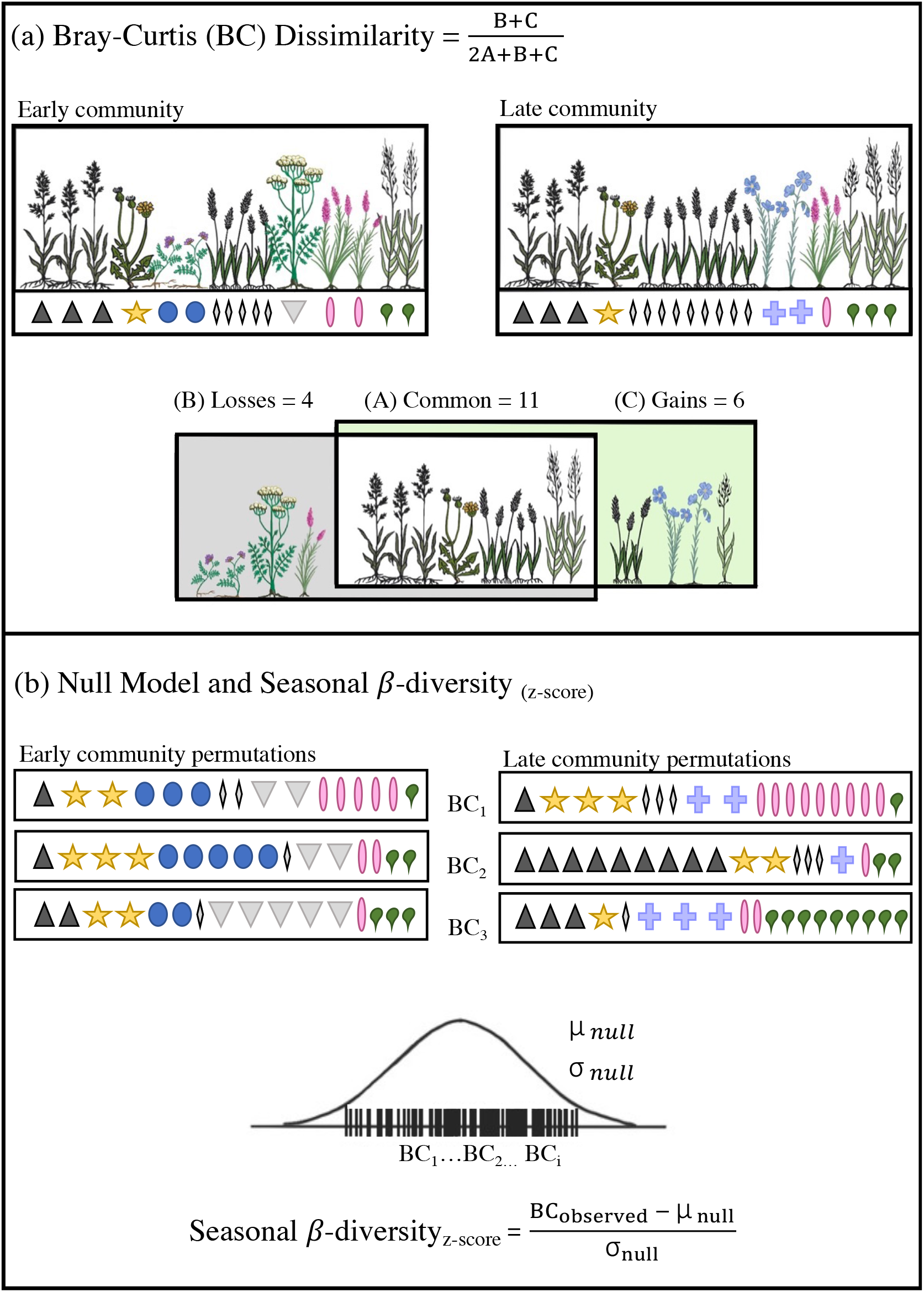
Panel a) Bray-Curtis dissimilarity is calculated using the formula 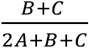 where B denotes species lost from the early to late community, C denotes species gained from the early to late community, and A denotes shared species between the two communities. Panel b) Null expectations of dissimilarity are obtained by permuting species abundances n number of times (3 times shown) while holding richness and total abundance within communities constant and calculating Bray-Curtis dissimilarity from these permuted communities. These values provide a distribution from which null mean (*μ_null_*) and variance (*σ_null_*) values of dissimilarity are estimated. Deviation (i.e., Seasonal β-diversity (z-scores)) values are then obtained using the following formula: Seasonal β – diversity 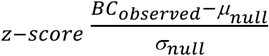 where BC_observed_ is the observed Bray-Curtis value from each early to late comparison, μ_null_ is the expected mean from the distribution of permutation comparisons, and σ_null_ is the variance from the distribution of permutation comparisons. Symbols in both panels represent species.

We also decomposed observed Bray-Curtis dissimilarity from each early-to-late comparison for each plot, year, and treatment into components of species “replacement” and “nestedness” (Baselga 2009). Nestedness captures the component of dissimilarity that results from one community being a subset of another community, whereas replacement captures the component of dissimilarity that results from certain species being lost and others gained between compared communities. We used the beta.part package (Baselga & Orme 2012) to obtain all dissimilarity estimates.

### Statistical models

We used two separate linear regression models to assess relationships between intra-annual temperature and precipitation variability and seasonal β-diversity of plant communities. In these models, average intra-annual temperature variability or precipitation variability for each site were included as a continuous predictor and average seasonal β-diversity estimates from control treatments from each site were included as a continuous response. We examined how NPKμ treatment influenced seasonal β-diversity using a multilevel regression model with plot-level seasonal β-diversity values included as response variables, treatment as a predictor, and a random effect of block, nested within treatment year, nested within site. Because not all functional groups are present at all sites, functional group analyses were completed using data from sites that had specific functional groups present at >1% cover overall. We fit five separate multilevel regression models to quantify how abundance of specific functional groups (i.e., summed cover of all species belonging to annual forbs, perennial forbs, C3 grasses, C4 grasses and legumes from m^2^ plots) differed among early and late sampling timepoints and treatments. These models included treatment by season combinations as categorical predictors (i.e., control_early, control_late, NPK*μ*_early, NPK*μ*__late) and block nested within treatment year nested within site as a random effect (i.e., interactions were not assessed prior to investigating differences in means).

For Bayesian inferences and estimates of uncertainty, all models described were fitted using the Hamiltonian Monte Carlo (HMC) sampler using Stan (Carpenter et al. 2017) and coded using the ‘brms’ package (Bürkner et al. 2018) in R (version V.2.1 R Core Development Team). All models were fit with 4 chains and 3000 iterations (Table S2). We used default priors for all models in which dissimilarly estimates (i.e., seasonal β-diversity, nestedness, turnover) were included as response variables. In models assessing differences in functional group abundances between timepoints and treatments, default priors were used for C3 and C4 graminoid and perennial forb models, whereas weakly regularizing priors were used in models assessing differences in legume and annual forb abundances (Table S2). We inspected the HMC chains to assess model convergence. For all models, we estimated the significance of effects by computing the difference between posterior distributions of interest and assessing whether the 90% credible interval of the difference contained zero.

## Results

We found a positive relationship between intra-annual temperature variability and seasonal β-diversity in unfertilized control plots (Slope: 0.0028, 90% Credible Interval (CI): 0.0004 to 0.0050) (Figure 2a) and no relationship between intra-annual precipitation variability and seasonal β-diversity (Slope: −0.0052, CI: −0.0278 to 0.0166) (Figure 2b). Contrary to our hypothesis, we detected a significant positive effect of fertilization on seasonal β-diversity across sites (Figure 3). This result indicates greater dissimilarity between early and late season communities in fertilized conditions, even after accounting for differences in richness between the two treatments (average number of species per m^2^ in control: 12.8; average number of species per m^2^ in NPK*μ*: 11.5). However, seasonal β-diversity values were negative in most cases, indicating lower than expected dissimilarity between early-season and late-season communities. After decomposing observed seasonal β-diversity into components of nestedness and turnover, we found no differences between treatments in nestedness (Figure 4a) but higher turnover in fertilized conditions than in controls (Figure 4b).

**Figure 2:**
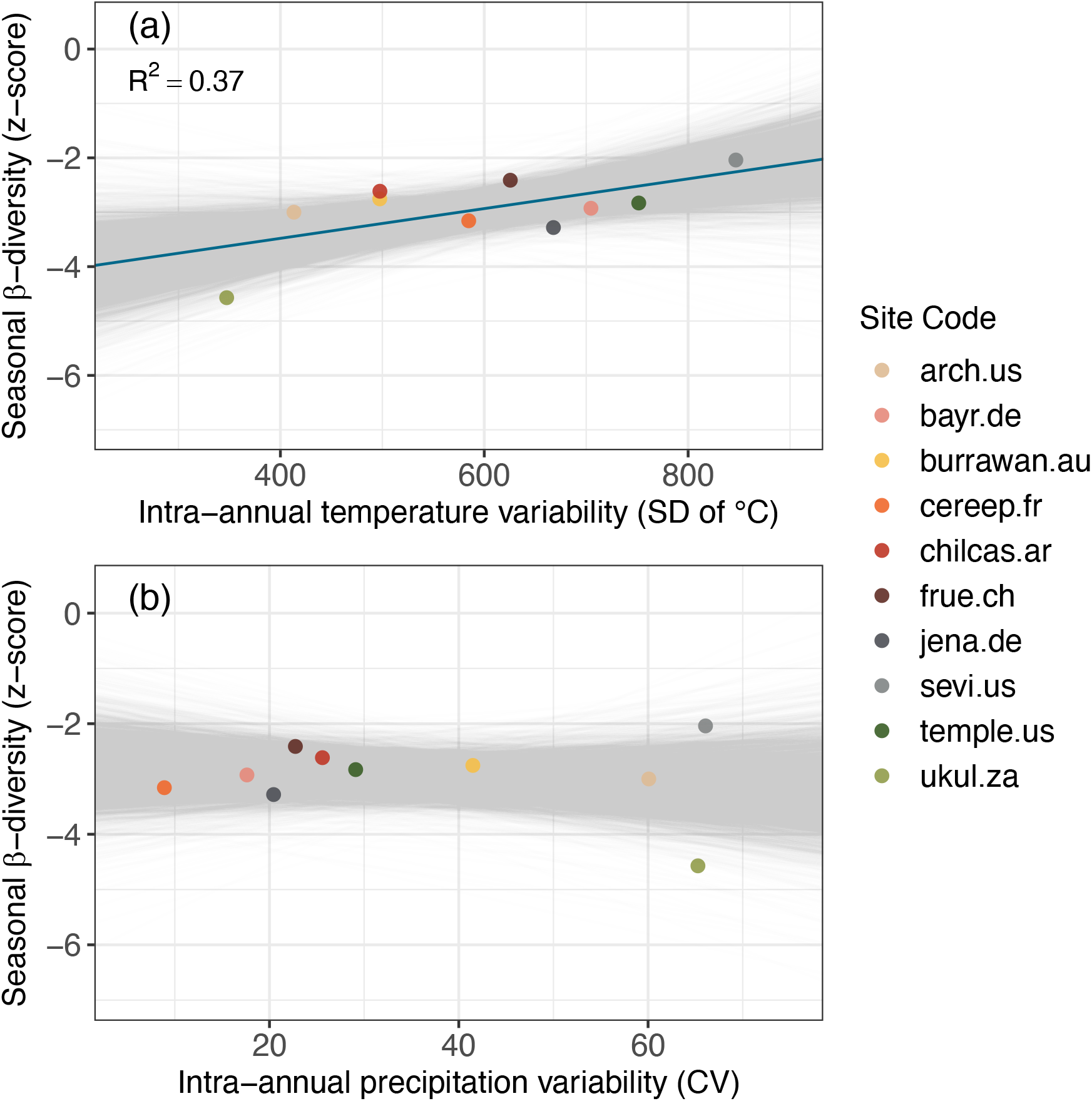
Relationships between intra-annual temperature (panel a) and precipitation (panel b) variability and mean site-level seasonal β-diversity in control treatments. Gray lines show 500 predicated relationships from posterior model estimates and dark blue lines show averages of these expectations when the distribution of the slope estimate differed from zero at the *α* = 0.1 probability level.

**Figure 3:**
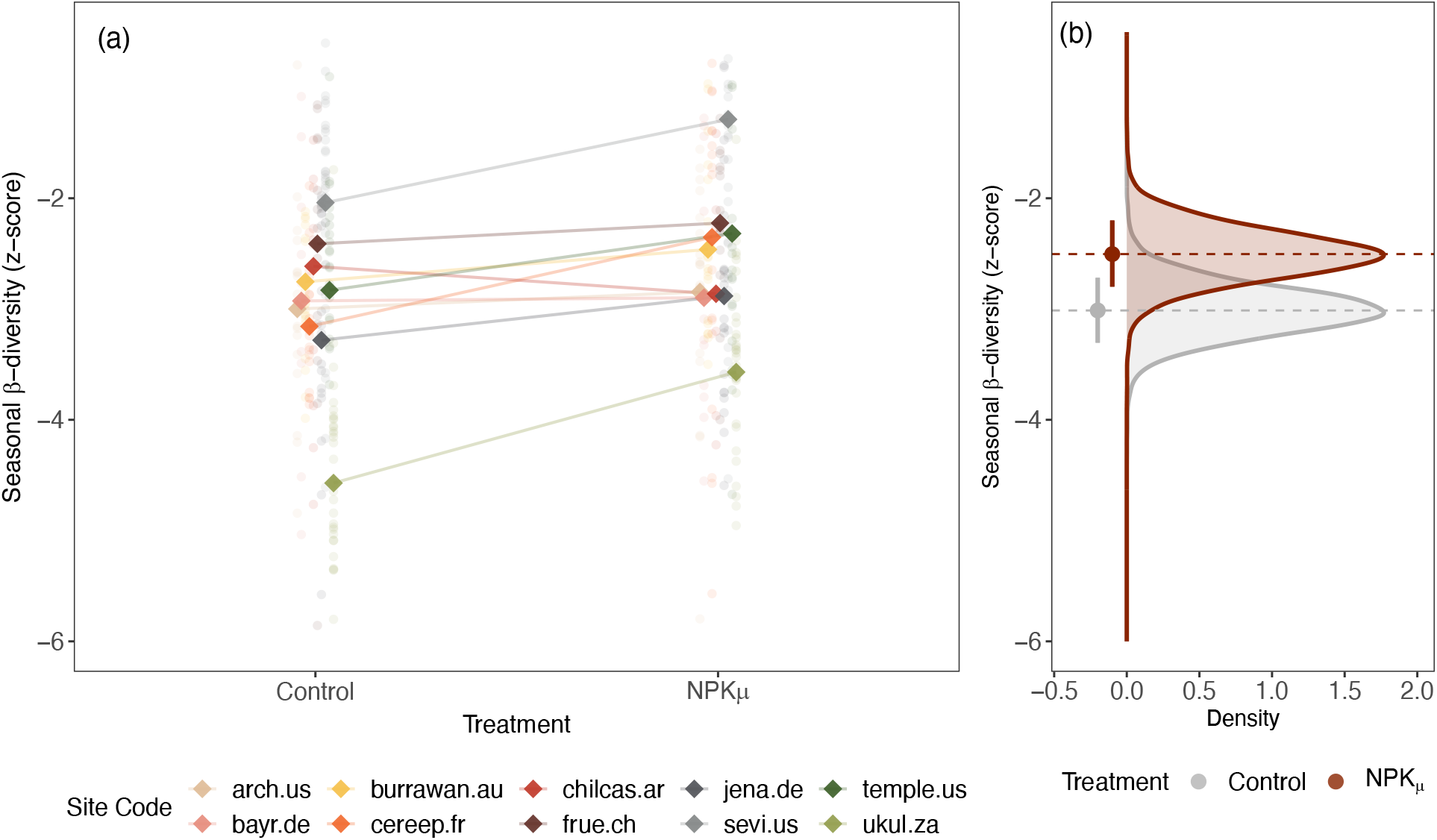
Panel a) Seasonal β-diversity in control and fertilized (i.e., NPKμ) treatments. Circle points are seasonal β-diversity values (i.e., z-scores) from each plot in each year for a given site (different colors) and diamond points are average site-level seasonal β-diversity estimates for each site in each treatment. Panel b) Posterior distributions of seasonal β-diversity in control and NPKμ treatments across all sites and years included in analyses. Horizontal lines and points denote estimated means and error bars denote 90% credible intervals of the distributions.

**Figure 4:**
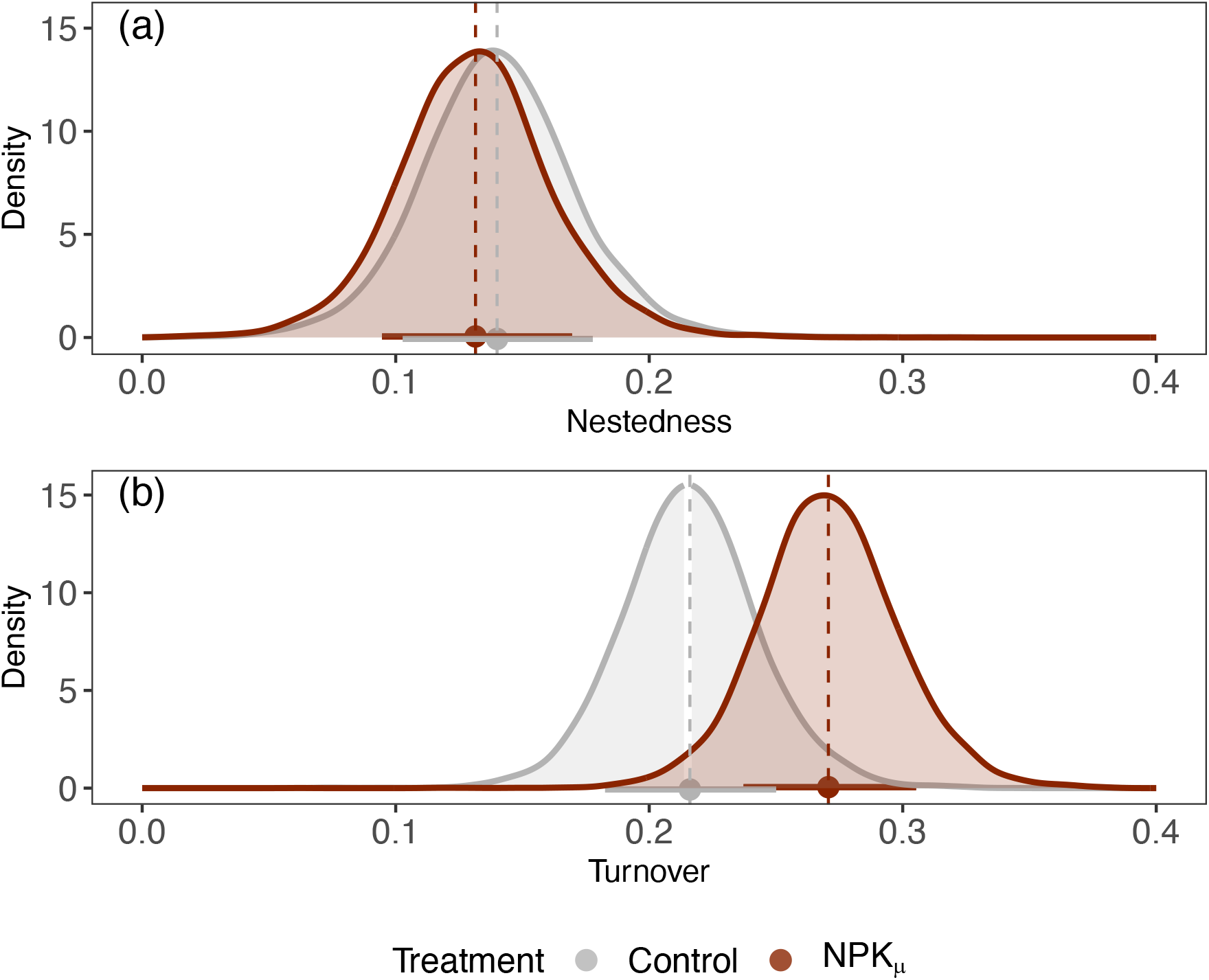
Posterior distributions of nestedness (panel a) and turnover (panel b) components of observed Bray-Curtis dissimilarity in control and fertilized (i.e., NPK) treatments across all sites and years included in analyses. Points and vertical lines denote estimated means and horizonal line segments denote 90% credible intervals of the distributions.

The effects of fertilization varied by functional group and between early and late sampling timepoints (Figure 5). At both sampling timepoints, fertilization resulted in higher cover of annual forbs (Figure 5a) and C3 graminoids (Figure 5c) and lower cover of C4 graminoids (Fig. 5d) and legumes (Fig. 5e). Perennial forb cover was reduced with fertilization, but only at the late sampling timepoint (Fig. 5b). Patterns of functional group abundances between early and late sampling points also varied between control and fertilized treatments. In control but not fertilized treatments, C4 graminoid abundance was higher at the later sampling point compared to early in the season and legume abundance was higher early in the season compared to late in the season. In contrast, abundance of C3 graminoids was higher early in the season compared to late in the season but only in fertilized treatments.

**Figure 5:**
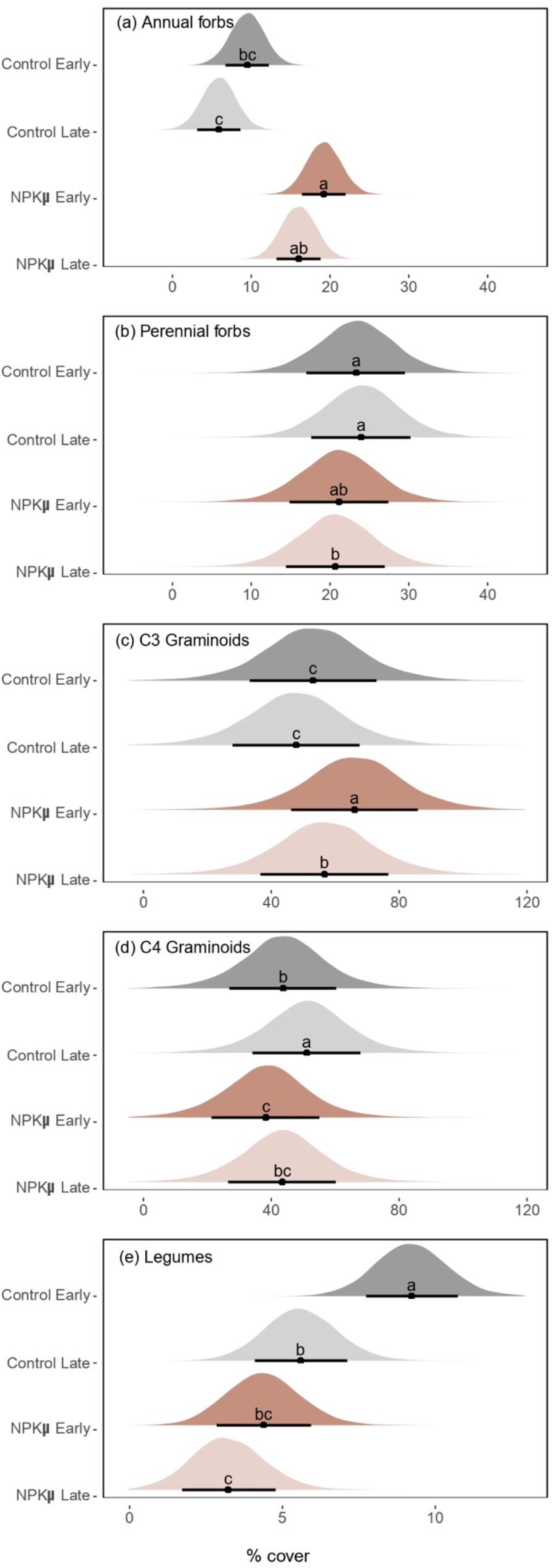
Posterior distributions of annual forbs (panel a, 6 sites included in analyses), perennial forbs (panel b, 10 sites included in analyses), C3 graminoids (panel c, 6 sites included in analyses), C4 graminoids (panel d; 6 sites included in analyses), and legumes (panel e; 8 sites included in analyses) early (dark colors) and late (light colors) in the growing season in control (grey) and fertilized (red) treatments. Different letters denote differences between groups at the *α* = 0.1 probability level, points denote mean estimates, and horizonal line segments denote 90% credible intervals of the distributions. Please note the different scales on the x-axes.

## Discussion

We found higher seasonal β-diversity in grasslands with high intra-annual temperature variability but observed no relationship between intra-annual precipitation variability and seasonal β-diversity. These finding suggests that positive relationships between α-diversity and intra-annual temperature variability identified in earlier studies (e.g., Scheiner & Rey-Benayas 1994, Boonman et al. 2021) may be underpinned by the positive influence of intra-annual temperature variability on temporal segregation of species within seasons. However, our findings also contradict several studies that implicate intra-annual precipitation variability as a key driver of seasonal β-diversity (e.g., Mulroy & Rundel 1977, Mathias & Chesson 2013). We suspect these contradictions arises due to differences in annual systems from which most evidence relating intra-annual precipitation variation to seasonal community dynamics has been gathered, and the predominantly perennial systems in our study. In annual systems, population, community, and ecosystem dynamics are often governed by interactions between intra-annual precipitation patterns and demographic traits related to dormancy and germination (e.g., Levine et al. 2011, Kimball et al. 2011, Shaw et al. 2022). In addition, in systems co-dominated by annuals and perennials, intra-annual variability in precipitation is often necessary for the persistence of annuals that occupy unique temporal niches in communities (e.g., Pérez-Camacho et al. 2012). On the other hand, intra-annual temperature variability may be a more important driver of intra-annual dynamics in perennial systems because temperature regulates plant physiology and shapes temporal segregation among many perennial plant species (e.g., Kemp & Williams. 1980, Monson et al. 1983). Given the low abundance of annual cover across our study sites (*c*. 11%), we are not able to test these patterns robustly. But a positive, non-significant, relationship between annual seasonal β-diversity and intra-annual precipitation variability (Supplementary Figure 1a) and a negative, non-significant, relationship between perennial seasonal β-diversity and intra-annual precipitation variability (Supplementary Figure 1b) invite new questions about how intra-annual precipitation variability influences seasonal community dynamics across diverse ecosystems.

The negative impacts of nutrient enrichment on grassland diversity have been extensively documented (e.g., Borer et al. 2017) and recent studies have identified a positive effect of fertilization on temporal β-diversity at inter-annual scales (Koerner et al. 2016, Hodapp et al. 2018, Chen et al. 2021). Mirroring these results at intra-annual scales, we found that fertilization resulted in higher seasonal β-diversity. Because we used deviations from null expectations as our measure of seasonal β-diversity, we can be confident that these shifts arise from true changes in composition rather than random processes influenced by richness. Similar to results found at inter-annual scales by Chen et al. 2021, we found that higher temporal β-diversity with fertilization was driven by higher turnover of species, and not nestedness of species, between early and late-season communities. This suggests that fertilization increases temporal β-diversity among and within years by similar mechanisms, namely by unique species occupying space within the community at different times (i.e., turnover) rather than species being lost from one timepoint to another (i.e., nestedness). At inter-annual scales, increased temporal β-diversity and species turnover through time can result in reduced stability of ecosystem productivity (e.g., Koerner et al. 2016, Chen et al. 2021). Additional research focused on how nutrient enrichment alters composition and associated ecosystem functions within growing seasons would clarify whether similar patterns manifest at intra-annual scales.

As expected, fertilization resulted in higher abundance of resource acquisitive species (i.e., C3 graminoids, annual forbs) and lower abundance of resource conservative species (i.e., C4 graminoids, legumes, perennial forbs) within and across sampling timepoints. Numerous other studies have documented similar patterns at single timepoints (e.g., Suding et al. 2005, Isabell et al. 2013, Wilcots et al. 2021, Tognetti et al. 2021) and results are primarily attributed to physiological differences between these groups and concomitant shifts from below-ground competition for nutrients to aboveground competition for light (e.g., Borer et al. 2014, Harpole et al. 2017). Among other anatomical and physiological differences, compared to C4 species, C3 species (i.e., most of the annuals in our study and C3 graminoids) generally have lower C:N ratios (e.g., Wedin & Tilman 1990), lower water and photosynthetic nitrogen use efficiencies (Black 1971, Taylor et al. 2011, Ripley et al. 2010), and earlier phenologies due to lower optimal temperatures for photosynthesis (Kemp & Williams 1980). These characteristics can lead to higher sensitivities of resource acquisitive species to nutrient limitation which allow them to increase in fertilized conditions (e.g., Zhong et al. 2019).

Fertilization made apparent seasonal differences in C3 graminoid abundance that were not present under control, while eliminating seasonal differences in abundance of C4 graminoids and legumes that were present in control treatments. Higher C3 graminoid abundance early in the season in fertilized treatments could reflect a higher capacity for growth of these species early in the season, when resources are abundant, followed by more pronounced declines later in the season when resources are scarce (e.g., Yuan et al. 2007). In contrast, both C4 graminoids and legumes are adapted to resource-limited conditions which can vary throughout growing seasons (Klaus et al. 2014). By altering resource conditions, fertilization likely modulates the low-resource periods during which these species thrive resulting in lower abundances of these species overall and homogenization of these groups across the growing season.

Our understanding of how grasslands respond to various components of global change is primarily based on studies that document community shifts at inter-annual scales. However, global changes may also disrupt within-season community dynamics. Here we show that seasonal β-diversity across 10 global grasslands is shaped by intra-annual temperature variability and eutrophication. Given that both intra-annual temperature variability (e.g., Xu et al. 2013) and precipitation variability (e.g., Hajek & Knapp 2021) are expected to increase in coming decades understanding effects of these understudied components of global change is essential to understanding how plant communities will shift into the future. Investigations focused on specific components of climatic variability such as seasonality (i.e., the occurrence of certain events within a definite limited period, *sensu* Lieth 1974) or predictability (i.e., the regularity of recurrence of the within cycle distribution of events, *sensu* Tonkin 2017) would further illuminate how fluctuations in climate shape seasonal dynamics of plant communities. We also show that nutrient enrichment increases seasonal β-diversity and species turnover, enhances the abundance of resource-acquisitive species (i.e., annual forbs, C3 graminoids) within and between timepoints, and decreases abundances of resource-conservative species (i.e., C4 graminoids, legumes) within and between timepoints. In addition, our results suggest that fertilization results in homogenization of abundances of resource conservative species early to late in the growing season. If the effects of intra-annual community change mirror those at inter-annual scales, the community shifts we observed in our study could have cascading impacts on ecosystem stability, multifunctionality, and the ability of these systems to recover from future perturbations. Our study provides new insight into the mechanisms by which climate variability and nutrient enrichment shape within-season community dynamics in global grasslands and highlights how discerning these patterns is essential to our understanding of biodiversity in these valuable ecosystems.

## Acknowledgements

Funding for this project was provided by the German Science Foundation (DFG; via University Leipzig/iDiv) for the project “Strategisches Projekt: NEu - Nutrient Network Europe” (Projektnummer 346001057-03). The authors do not have any conflicts of interest to declare.

## Author contributions

MG and SH conceptualized this project, MG completed analyses with assistance from HH, SJ, and EL, MG wrote the manuscript, all authors contributed to data collection and editing the manuscript.

