## Supplementary Information for "Intra-annual temperature variability and nutrient enrichment drive seasonal β-diversity in global grasslands"

Supplementary Information to the paper “Temperature seasonality and nutrient enrichment drive intra-annual community turnover in global grasslands”

Table S1: Nutrient Network sites from which compositional data were used in this study

| Site Code | Country | Latitude | Longitude | Years of data |
| --- | --- | --- | --- | --- |
| arch.us | United States | 27.18 | -81.35 | 6 |
| bayr.de | Germany | 49.91 | 11.58 | 4 |
| burrawan.au | Australia | -27.73 | 151.14 | 7 |
| cereep.fr | France | 48.29 | 2.68 | 6 |
| chilcas.ar | Argentina | -36.28 | -58.27 | 4 |
| frue.ch | Switzerland | 47.11 | 8.54 | 5 |
| jena.de | Germany | 50.95 | 11.62 | 7 |
| temple.us | United States | 31.04 | -97.35 | 8 |
| sevi.us | United States | 34.36 | -106.69 | 7 |
| ukul.za | South Africa | -29.67 | 30.40 | 11 |

Table S2: All models were fit using the brms package in R (Bürkner 2022) with 2,000 iterations for warmup. We visually inspected predicted vs. observed values and distributions of residuals to assess model performance. For functional group abundance models, we included block nested within treatment year nested within site as a random effect. “trt_sampling” refers to categorical variables created for treatment and sampling timepoint combinations (i.e., control_early, control_late, NPK$\mu$_early, NPK$\mu$_late).

| Model | Distributions | Iterations | Effective Sample Sizes | Rhat |
| --- | --- | --- | --- | --- |
| Climate variability models | | | | |
| Seasonal β ~ intra-annual temperature variability | Gaussian | 3,000 | 1,793 - 3,346 | 1 |
| Seasonal β ~ intra-annual precipitation variability | Gaussian | 3,000 | 2,736 - 2,131 | 1 |
| Treatment models | | | | |
| Seasonal β ~ trt | Gaussian | 3,000 | 2,881 - 8620 | 1 |
| nestedness ~ trt | Gaussian | 3,000 | 2,181 - 3,044 | 1 |
| turnover ~ trt | Gaussian | 3,000 | 2,191 - 2,769 | 1 |
| Functional group abundance models | | | | |
| Annual forbs ~ trt_sampling | Gaussian, Intercept: Gaussian (12, 2) | 3,000 | 2,717 - 3,949 | 1 |
| Perennial forbs ~ trt_sampling | Gaussian | 3,000 | 1,153 - 4,915 | 1 |
| C3 graminoids ~ trt_sampling | Gaussian | 3,000 | 1,226 - 4,863 | 1 |
| C4 graminoids ~ trt_sampling | Gaussian | 3,000 | 1,129 - 4,583 | 1 |
| Legumes ~ trt_sampling | Gaussian, Intercept: Gaussian (8, 2) | 3,000 | 3,444 - 6,138 | 1 |

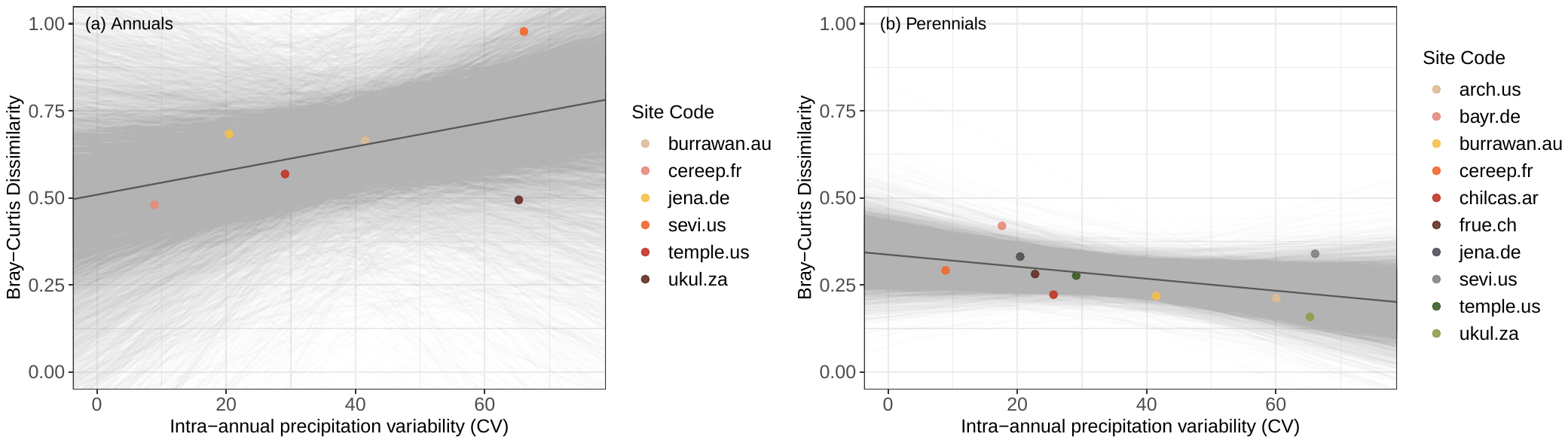

Figure S1: Relationships between precipitation seasonality and Bray-Curtis dissimilarity of annual (panel a) and perennial (panel b) species across study sites in which they occur. Neither relationship is significant at the $\alpha$ = 0.1 level.
